# Dysregulation of hippocampal adult-born immature neurons disrupts a brain-wide network for spatial memory

**DOI:** 10.1101/2020.09.02.273649

**Authors:** Hechen Bao, Zhiqiang Hu, Sung-ho Lee, Ramya Kolagani, Tzu-Hao Harry Chao, Yan-Jia Luo, Woomi Ban, Heather Anne Sullivan, Sergio Gamero-Alameda, Zane R. Lybrand, Yuguo Yu, Jenny Hsieh, Ian R Wickersham, Steven E Brenner, Yen-Yu Ian Shih, Juan Song

## Abstract

Mounting evidence suggests that cognitive deficits associated with various neurological disorders may arise in part from a small population of dysregulated adult-born neurons in the dentate gyrus (DG). How these dysregulated adult-born neurons contribute to brain-wide network maladaptation and subsequent cognitive deficits remains unknown. Using an established mouse model with a small number of time-stamped dysregulated adult-born immature neurons and spatial memory deficits, we performed resting state functional magnetic resonance imaging and found that approximately 500 deficient immature neurons (<0.1% of total DG granule neurons) are sufficient to induce a significant decrease in the functional connectivity between DG and insular cortex (IC), two brain regions without direct anatomical connections. Furthermore, using a combination of rabies-based retrograde tracing and *in vivo* fiber photometry recording, we demonstrated that dysregulated adult-born neurons induce aberrant activity and synchrony in local hippocampal CA3 and CA1 regions, as well as distal medial-dorsal thalamus and IC regions during a spatial memory process. These results suggest that a few hundred dysregulated adult-born immature neurons can impact brain-wide network dynamics across several anatomically discrete regions and collectively contribute to impaired cognitive functions.

## Introduction

Within adult hippocampus, the dentate gyrus (DG) continuously generates new granule neurons throughout life in almost all mammals, including humans (Boldrini et al., 2018; Ming and Song, 2011; Moreno-Jimenez et al., 2019; Spalding et al., 2013; Zhao et al., 2008). In the past decade, adult hippocampal neurogenesis has garnered significant interests because of its proposed role as a neural substrate for cognitive function and a pathophysiological site associated with various neurological disorders. Substantial evidence in rodent models has supported a causal role of adult-born immature neurons in specific forms of learning and memory, such as spatial memory and pattern separation (Akers et al., 2014; Clelland et al., 2009; Miller and Sahay, 2019; Sahay et al., 2011). In contrast, dysregulated adult-born immature neurons have been shown to contribute to the cognitive deficits associated with mental disorders and epilepsy (Bao and Song, 2018; Christian et al., 2014; Kang et al., 2016; Zhou et al., 2013). These studies suggest that adult-born immature neurons with unique physiological or pathological properties can potentially modulate local hippocampal circuitry and impact hippocampal dependent learning and memory. Supporting this view, recent studies have demonstrated that adult-born immature neurons with enhanced plasticity during a critical window are capable of modulating the activity of various cell types within the local hippocampal circuitry, including mature granule cells, hilar interneurons, and CA3 pyramidal cells (Alvarez et al., 2016; Gu et al., 2012; Ikrar et al., 2013; Temprana et al., 2015). However, a major gap remains in our understanding is whether and how a small population of time-stamped adult-born immature neurons with transient and special properties contribute to brain-wide alteration of the neural network associated with learning and memory, both within and beyond hippocampal formation.

To address this question, we took advantage of an established mouse model with a small number of time-stamped dysregulated adult-born immature neurons via retrovirus-mediated knockdown of *Disrupted-in-schizophrenia 1* (DISC1) gene, which exhibits hippocampal dependent spatial memory deficits (Zhou et al., 2013). Using a combination of fMRI, rabies-based retrograde tracing, *in vivo* multi-fiber photometry recording, and network analysis, we provided novel network-level evidence that ~500 dysregulated newborn neurons (<0.1% of total DG granule neurons) are sufficient to induce brain-wide maladaptation across multiple anatomically discrete regions during the spatial memory process, ranging from local hippocampal CA3 and CA1 regions to distal medial-dorsal thalamic (MDTH) and insular cortical (IC) regions. These results provide the first experimental evidence that a few hundred dysregulated adult-born immature neurons can impact brain-wide network dynamics across several anatomically discrete regions within and beyond the hippocampal formation to collectively impact memory.

## Results

### Dysregulated adult-born immature neurons disrupt DG-IC functional connectivity and IC activity during spatial memory behavior

To specifically target adult-born immature neurons, we delivered retroviruses co-expressing GFP and a previously validated short hairpin RNA (shRNA) against mouse DISC1 (shDISC1) or control shRNA (shControl) to the adult DG for birth-dating and genetic manipulation of newborn neurons (Fig. 1A), and assessed the impact of DISC1 deficiency on the morphology of newborn neurons and cognitive behavior of retroviral injected mice at 18 days post injection (dpi) that is within a time window behavioral deficits were observed (Zhou et al., 2013) (referred as DISC1 mice thereafter). Consistent with previous reports (Duan et al., 2007; Kang et al., 2011; Kim et al., 2012), we found a series of morphological deficits in DISC1-deficient newborn neurons, including soma hypertrophy, aberrant neuronal migration, and accelerated maturation (SFig. 1 AC). Moreover, DISC1-deficient newborn neurons exhibited increased intrinsic excitability (Fig. 1B, SFig. 1D). Importantly, DISC1 mice with approximately 500 retroviral labeled newborn neurons (<0.1% of total DG granule neurons) exhibited spatial memory deficits during a hippocampal dependent novel place recognition (NPR) test (Fig. 1C-E) without affecting the locomotor activity (SFig. 1E-F). These results suggest that deficiency in a small number of newborn neurons is sufficient to cause spatial memory deficits presumably by affecting neural circuits and networks associated with spatial memory functions.

**Figure 1.**
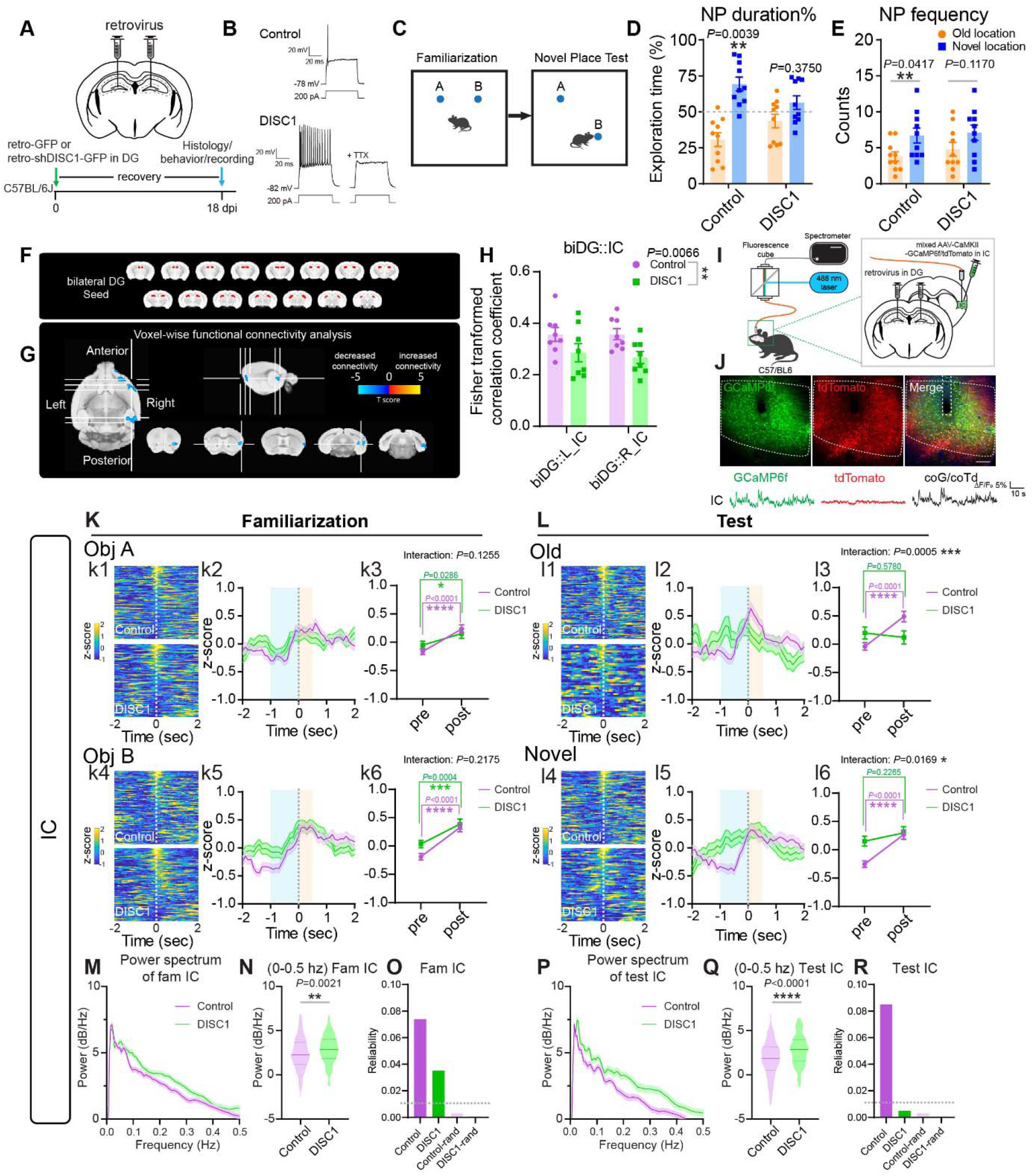
Dysregulated adult-born immature neurons disrupt DG-IC functional connectivity and IC activity during spatial memory behavior. (A) Experimental scheme of surgery and assays. (B) Sample voltage responses of adult-born neurons responding to a 500-ms current injection at 18 dpi. (C) Illustration of novel place recognition test paradigm. (D) Exploration duration% during test phase. (n=10 per group, Wilcoxon signed rank test (vs 50%), \*\**P*(control)=0.0039, *P*(DISC1)=0.3750). (E) Sniffing frequency during test phase. (n=10 per group, two-way RM ANOVA followed by Sidak’s *post hoc* test. \**P*(control)=0.0417, *P*(DISC1)=0.1170). (F) Representative images showing bilateral DG as the seeding region. (G) Voxel-wise analysis showing voxels that had altered connectivity with biDG were accumulated in the distal insular cortex. (thresholded at p < 0.05 and #voxels > 100) (H) Correlation coefficient comparison of biDG::IC between control and DISC1. (n=8 per group, two-way ANOVA to compare Control vs DISC1, *F*_1,28_=8.63, \*\**P*=0.0066) (I) Illustration of *in vivo* fiber photometry recording system. (J) Upper: Composite confocal image showing GCaMP6f and control tdTomato expression in IC. Scale bar, 100 μm. Lower: Representative signal traces. (K-L) Fiber photometry recording of IC during NPR. (k1,k4,l1,l4) Heatmaps of all trials of individual mice during familiarization or test events. (k2,k5,l2,l5) Mean responses of all trials during events. (k3,k6,l3,l6) Quantification of z-score comparisons of pre-(−1~0s) and post-(0~0.5s) event. Statistics were performed by two-way RM ANOVA followed by Holm-Sidak’s *post hoc* test. Fam (Object A: n(control/DISC1)=141/108 trials, main effect of Interaction *F*_1,115_=2.364, *P*=0.1255, \*\*\*\**P*(control)<0.0001, \**P*(DISC1)=0.0286. Object B: n(control/DISC1)=123/106 trials, main effect of Interaction *F*_1,227_=1.529, *P*=0.2175, \*\*\*\**P*(control)<0.0001, \*\*\**P*(DISC1)=0.0004). Test (Old object: n(control/DISC1)=91/46 trials, main effect of Interaction *F*_1,135_=12.53, \*\*\**P*=0.0005, \*\*\*\**P*(control)<0.0001, *P*(DISC1)=0.5780. Novel object: main effect of Interaction *F*_1,183_=5.817, \**P*=0.0169, n(control/DISC1)=113/72 trials, \*\*\*\**P*(control)<0.0001, *P*(DISC1)=0.2265). All trials were from 8 mice per group. Dashed lines indicate onset of the events. \**P*<0.05, \*\**P*< 0.01, \*\*\**P*<0.001, \*\*\*\**P*<0.0001. (M) Power spectral density of IC during familiarization. (N) Comparison of averaged power of 0-0.5Hz in IC during familiarization. (n(control/DISC1)=264/214 trials from 8 mice per group. Two-tailed unpaired Student’s *t* test, \*\**P*=0.0021). (O) Reliability of IC Ca^2+^ responding to events during familiarization (n(control/DISC1)=264/214 trials, n(control-rand/DISC1-rand)=240/240 non-event trials were randomly selected from 8 mice per group). Detailed in Methods. (P) Power spectral density of IC during test. (Q) Comparison of averaged power of 0-0.5Hz in IC during test. (n(control/DISC1)=204/118 trials from 8 mice per group. Two-tailed unpaired Student’s *t* test, \*\*\*\**P*<0.0001). (R) Reliability of IC Ca^2+^ responding to events during test (n(control/DISC1)=204/118 trials, n(control-rand/DISC1-rand)=240/240 non-event trials were randomly selected from 8 mice per group). Dashed lines in (O,R) indicate thresholding reliability at 0.01. Values represent mean ± SEM.

Next, we sought to address whether dysregulated newborn neurons contribute to brainwide network maladaptation in DISC1 mice. As an entry point, we took a brain-wide approach by performing resting state functional magnetic resonance imaging (rsfMRI). After preprocessing of the rsfMRI data, we generated voxel mask of bilateral DG (biDG) and preformed seed-based voxel-wise analysis (Fig. 1F). This approach allows us to examine the whole brain at the voxel level to detect regions of interest (ROIs) that exhibit altered correlation with biDG in DISC1 mice (Gorges et al., 2017). Interestingly, we found that a cluster of voxels in insular cortex (IC) exhibit decreased correlation with biDG in DISC1 mice as compared to the control mice (Fig. 1G). Furthermore, we defined ROIs at IC and DG based on anatomical atlas and examined the extent of IC-DG correlation. Consistent with the results obtained from seed-based voxel-wise analysis, we found that the correlation between biDG and IC significantly decreased in DISC1 mice (Fig. 1H). These results suggest that a small number of dysregulated newborn neurons in the DG are sufficient to induce altered functional connectivity with a distal cortical brain region.

The rsfMRI was performed in anesthetized mice, therefore, we sought to validate these findings in behaving mice. We hypothesized that decreased DG-IC correlation may result from altered IC activity during hippocampal dependent behavior. To test this, we utilized a spectrometer based in vivo fiber photometry system (Li et al., 2020; Meng et al., 2018) to record the excitatory neurons in the right IC labeled with AAV-CaMKII-GCaMP6f in the home cage and during the NPR test. AAV-CAG-tdTomato was co-injected to control the motion artifacts (Fig. 1I-J, SFig. 1G-H). The Ca^2+^ activity measured by the frequency and amplitude of the Ca^2+^ events (Beier et al., 2017) and the power spectral density(Tibau et al., 2013) of the dominant frequency component (PDF) (0-0.5 Hz) of the Ca^2+^ signals in the IC of DISC1 mice did not significantly alter in the home cage, as compared to control mice (SFig. 1N-R). However, during the NPR test, DISC1 mice showed a significant increase in the PDF of the Ca^2+^ signals in the IC during both familiarization and test phases, indicating that deficient newborn neurons altered IC neuronal firing pattern and traits during spatial memory (Fig. 1M-N, P-Q). Validation of these parameters was performed with negative controls including imaging non-Ca^2+^ activity dependent reporter or in the absence of reporter (SFig. 1I-M). Furthermore, we performed peri-event analysis (Calipari et al., 2016) by examining the GCaMP signals in control and DISC1 mice when they were e xploring the objects at distinct locations. We found that control mice showed a robust elevation of IC Ca^2+^ activity when they were exploring the objects/locations during both familiarization (memory encoding) and test (memory retrieval) phases (Fig. 1K-L). In contrast, despite DISC1 mice responding to objects/locations with a moderate elevation of IC Ca^2+^ activity during familiarization (Fig. 1K), they failed to respond to both object/locations with a robust elevation of IC Ca^2+^ signals during the test phase as compared to control mice (Fig. 1L). Consistent with these findings, control mice showed robust reliability (Haider et al., 2010), which measured the likelihood of Ca^2+^ dynamics in correlation with objects/locations, at both phases, while DISC1 mice showed a large decrease (53% less than control) in reliability during familiarization and no obvious reliability (lower than the threshold of 0.01) was observed during the test (Fig. 1O, R). Together, these data suggest that a small number of dysregulated newborn neurons are sufficient to induce aberrant IC activity during the test (but not the familiarization) phase of the spatial memory process.

### Dysregulated adult-born immature neurons disrupt activity of medial dorsal thalamus at baseline and during spatial memory behavior

The IC is known as an integration hub that contributes to various functions including cognition (Gogolla, 2017). However, the DG and IC are remote from each other anatomically. Therefore, we wondered whether the DG directly or indirectly connects with IC. To address this, we performed rabies-based retrograde tracing to map the direct monosynaptic inputs to IC (Fig. 2A). As a result, we identified several input regions to IC excitatory neurons, including contralateral somatosensory cortex (cS1), contralateral insular cortex (cIC), thalamus (TH), hypothalamus (HYTH), entorhinal cortex (EC), and amygdala (Amy) (Fig. 2B-L, SFig. 2A-B). Importantly, we did not identify the direct inputs from DG or hippocampus to IC (Fig. 2E-G, K), suggesting an indirect anatomical connection between DG and IC. Interestingly, among these inputs, the medial-dorsal thalamus (MDTH) exhibited high anatomical connectivity with IC excitatory neurons (Fig. 2H, L). Based on this high MDTH-IC connectivity, we sought to record the Ca^2+^ activity of excitatory neurons in MDTH from control and DISC1 mice in freely moving animals in the home cage and during the NPR test using in vivo fiber photometry (Fig. 2M, SFig. 2C). Interestingly, unlike the IC Ca^2+^ activity that showed no change between control and DISC1 mice in the home cage, we found that DISC1 mice exhibit a significant decrease in MDTH Ca^2+^ activity as compared to control mice without altering the peak amplitude Ca^2+^ events (Fig. 2N-P). The PDF of the Ca^2+^ events in the home cage or during the NPR test showed no significant differences (SFig. 3D-I). Interestingly, similar to the IC responses during the NPR test, control mice showed a robust elevation of MDTH Ca^2+^ activity during both familiarization and test phases. In contrast, DISC1 mice failed to respond to the objects/locations with a robust elevation of MDTH Ca^2+^ signals during both phases as compared to control mice (Fig. 2S-T). Moreover, control mice showed robust reliability of peri-event associated Ca^2+^ dynamics at both phases, while DISC1 mice showed a large decrease in reliability (63% less than control) during familiarization and no obvious reliability during the test (Fig. 2Q, R). Together, these data suggest that a small number of dysregulated adult-born neurons are sufficient to induce aberrant MDTH activity during the spatial memory process. Furthermore, the high similarity of the responses between IC and MDTH during the NPR test not only supports the high anatomical connectivity between these two brain regions, but also suggests that activities of IC and MDTH maybe highly correlated during hippocampal dependent spatial memory process.

**Figure 2.**
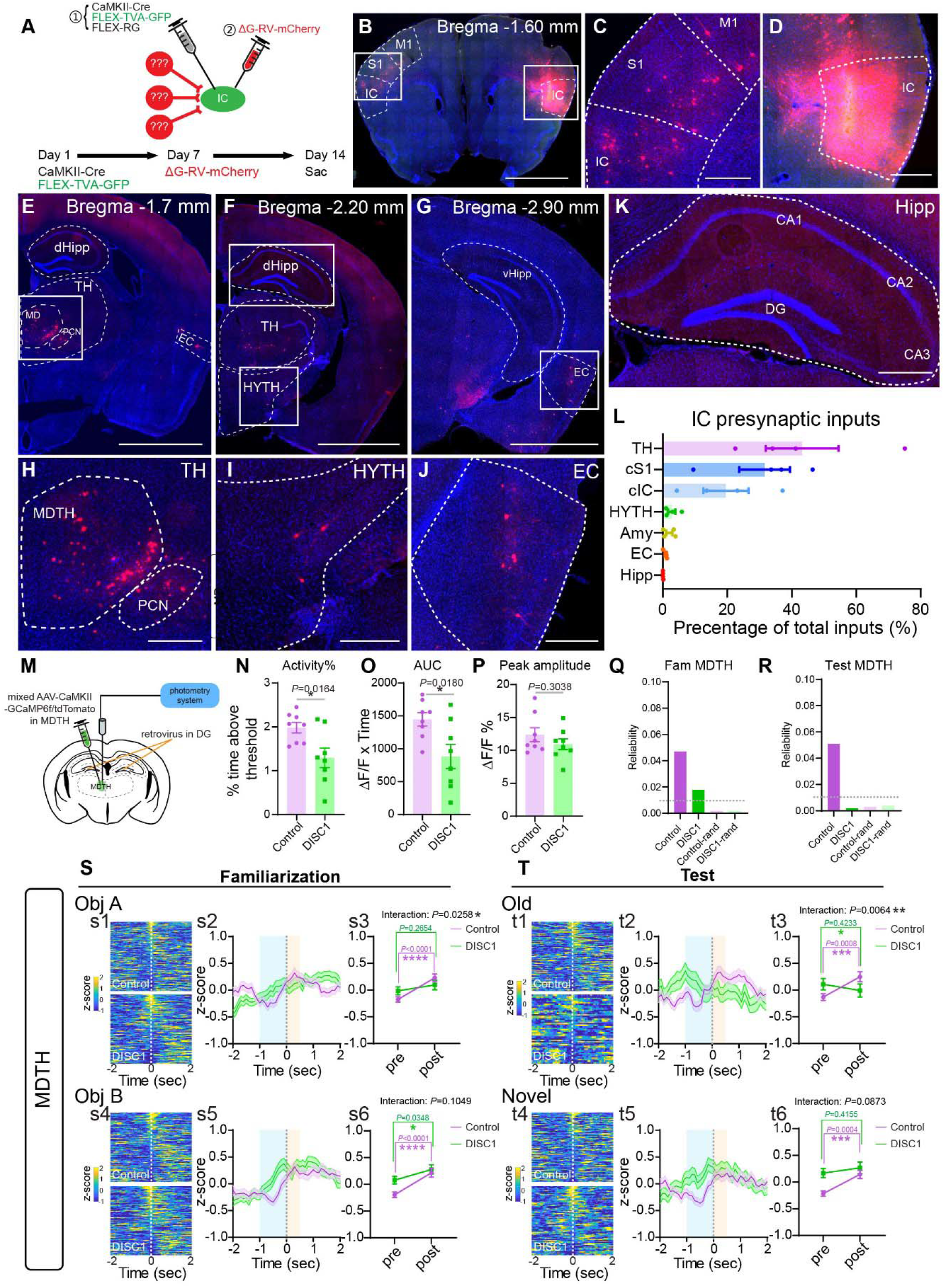
Dysregulated adult-born immature neurons disrupt activity of medial dorsal thalamus at baseline and during spatial memory behavior. (A) Experimental scheme of monosynaptic rabies retrograde tracing of IC. (B-D) Images of injection site and contralateral cortical inputs to IC. Scale bar, (B) 2 mm, (C, D) 400 μm. (E-G) Images of distal inputs from TH, HYTH and EC. Scale bar, 2 mm. (H-J) Zoomed images of TH, HYTH and EC inputs. Scale bar, 400 μm. (K) Image of hippocampus showing no direct hippocampal inputs to IC. Scale bar, 400 μm. (L) Percentage summary of IC inputs. (n=4 mice). (M) Experimental scheme of fiber photometry recording from MDTH. (N-P) MDTH homecage Ca^2+^ activity. (N) The percentage of high Ca^2+^ activity events among total recording session, \**P*=0.0164. (O) Area under the curve of high Ca^2+^ activity events, \**P*=0.0180. (P) Average peak amplitude of high Ca^2+^ activity event, *P*=0.3038. Two-tailed unpaired Student’s *t* test. n=8 mice per group. (Q) Reliability of MDTH Ca^2+^ responding to events during familiarization (n(control/DISC1)=264/214 trials, n(control-rand/DISC1-rand)=240/240 non-event trials were randomly selected from 8 mice per group). (R) Reliability of MDTH Ca^2+^ responding to events during test (n(control/DISC1)=204/118 trials, n(control-rand/DISC1-rand)=240/240 non-event trials were randomly selected from 8 mice per group). Dashed lines indicate thresholding reliability at 0.01. (S-T) Fiber photometry recording of MDTH during NPR. (s1,s4,t1,t4) Heatmaps of all trials of individual mice during familiarization or test events. (s2,s5,t2,t5) Mean responses of all trials during events. (s3,s6,t3,t6) Quantification of z-score comparisons of pre-(−1~0s) and post-(0~0.5s) event. Statistics were performed by two-way RM ANOVA followed by Holm-Sidak’s *post hoc* test. Fam (Object A: n(control/DISC1)=141/108 trials, main effect of Interaction *F*_1,247_=5.027, \**P*=0.0258, \*\*\*\**P*(control)<0.0001, *P*(DISC1)=0.2654. Object B: n(control/DISC1)=123/106 trials, main effect of Interaction *F*_1,227_=2.650, *P*=0.1049, \*\*\*\**P*(control)<0.0001, * *P*(DISC1)=0.0348). Test (Old object: n(control/DISC1)=91/46 trials, main effect of Interaction *F*_1,135_=7.666, \*\*\**P*=0.0064, \*\*\**P*(control)=0.0008, *P*(DISC1)=0.4233. Novel object: main effect of Interaction *F*_1,183_=2.954, *P*=0.0873, n(control/DISC1)=113/72 trials, \*\*\**P*(control)=0.0004, *P*(DISC1)=0.4155). All trials were from 8 mice per group. Dashed lines indicate onset of the events. \**P*<0.05, \*\**P*< 0.01, \*\*\**P*<0.001, \*\*\*\**P*<0.0001. Values represent mean ± SEM. Contralateral somatosensory cortex (cS1), contralateral insular cortex (cIC), thalamus (TH), medial dorsal thalamus (MDTH), paracentral thalamic nucleus (PCN), (HYTH), entorhinal cortex (EC), dorsal hippocampus (dHipp), ventral hippocampus (vHipp).

### Dysregulated adult-born immature neurons disrupt local hippocampal activity during spatial memory behavior

Having identified altered MDTH and IC activity resulted from dysregulated newborn neurons, we wondered whether local hippocampal regions could serve as intermediates to couple aberrant activity from dysregulated newborn neurons to these distal brain regions, including the MDTH and IC. Adult-born new neurons reside in a tri-synaptic circuit, and it was thought that these new neurons are capable of impacting the activity of their downstream hippocampal regions, such as CA3 and CA1 (Bao and Song, 2018; Christian et al., 2014; Tran et al., 2019). Therefore, we asked whether hippocampal CA3 or CA1 is a direct input region to MDTH (Fig. 3A). To address this question, we performed rabies-based retrograde tracing (Fig. 3B-C), and found that CA1 (but not CA3) pyramidal cells form direct synaptic connections with MDTH (Fig. 3D-E, SFig. 3A, M), suggesting that CA1 may serve as a gateway to relay information from the DG newborn neurons to the MDTH and then IC. Given that CA3 is the immediate downstream region of DG newborn neurons, we hypothesized that the information flow from dysregulated newborn neurons follows the CA3-CA1-MDTH-IC pathway. To test this hypothesis, we sought to address whether activity of the CA3 and CA1 alters in DISC1 mice at baseline and during behavior. We performed in vivo fiber photometry recording in CA3 and CA1 pyramidal cells of control and DISC1 mice in the home cage and during the NPR test (SFig. 3B-C, N-O). Interestingly, we found no change in the Ca^2+^ activity and PDF in both CA3 and CA1 between control and DISC1 mice in the home cage (SFig. 3D-L, P-X).

**Figure 3.**
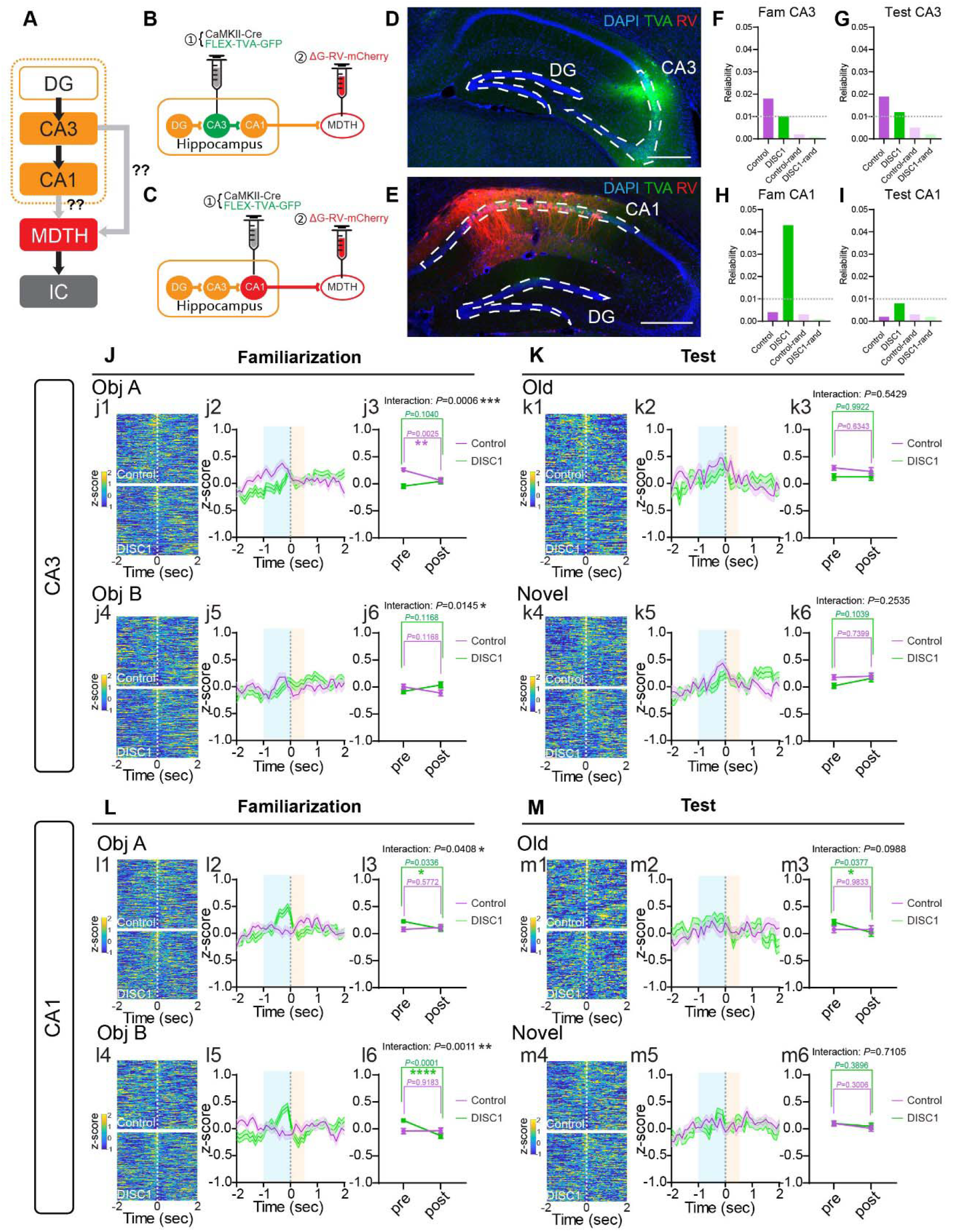
Dysregulated adult-born immature neurons disrupt local hippocampal activity during spatial memory behavior. (A) Illustration of circuit model. (B-C) Experimental scheme of monosynaptic rabies retrograde tracing of MDTH. (D) Sample image showing no direct inputs from CA3 to MDTH. Scale bar, 400 μm. n= 3 mice. (E) Sample image showing direct inputs from CA1 to MDTH. n=3 mice. Scale bar, 400 μm. (F) Reliability of CA3 Ca^2+^ responding to events during familiarization (n(control/DISC1)=303/353 trials, n(control-rand/DISC1-rand) =210/210 non-event trials were randomly selected from 7 mice per group). (G) Reliability of CA3 Ca^2+^ responding to events during test (n(control/DISC1)=200/198 trials, n(control-rand/DISC1-rand)=210/210 non-event trials were randomly selected from 7 mice per group). Dashed lines indicate thresholding reliability at 0.01. (H) Reliability of CA1 Ca^2+^ responding to events during familiarization (n(control/DISC1)=303/353 trials, n(control-rand/DISC1-rand)=210/210 non-event trials were randomly selected from 7 mice per group). (I) Reliability of CA1 Ca^2+^ responding to events during test (n(control/DISC1)=200/198 trials, n(control-rand/DISC1-rand)=210/210 non-event trials were randomly selected from 7 mice per group). Dashed lines indicate thresholding reliability at 0.01. (J-K) Fiber photometry recording of CA3 during NPR. (j1,j4,k1,k4) Heatmaps of all trials of individual mice during familiarization or test events. (j2,j5,k2,k5) Mean responses of all trials during events. (j3,j6,k3,k6) Quantification of z-score comparisons of pre-(−1~0s) and post-(0~0.5s) event. Statistics were performed by Two-way RM ANOVA followed by Holm-Sidak’s *post hoc* test. Fam (Object A: n(control/DISC1)=163/183 trials, main effect of Interaction *F*_1,344_=12.13, \*\*\**P*=0.0006, \*\**P*(control)=0.0025, *P*(DISC1)=0.1040. Object B: n(control/DISC1)=140/170 trials, main effect of Interaction *F*_1,308_=6.043, \**P*=0.0145, *P*(control)=0.1168, *P*(DISC1)=0.1168). Test (Old object: n(control/DISC1)=81/81 trials, main effect of Interaction *F*_1,160_=0.3717, *P*=0.5429, *P*(control)=0.6343, *P*(DISC1)=0.9922. Novel object: main effect of Interaction *F*_1,234_=1.310, *P*=0.2535, n(control/DISC1)=119/117 trials, *P*(control)=0.7399, *P*(DISC1)=0.1039). (L-M) Fiber photometry recording of CA1 during NPR. (l1,l4,m1,m4) Heatmaps of all trials of individual mice during familiarization or test events. (l2,l5,m2,m5) Mean responses of all trials during events. (l3,l6,m3,m6) Quantification of z-score comparisons of pre-(−1~0s) and post-(0~0.5s) event. Statistics were performed by two-way RM ANOVA followed by Holm-Sidak’s *post hoc* test. Fam (Object A: n(control/DISC1)=163/183 trials, main effect of Interaction *F*_1,344_=4.21, \**P*=0.0408, *P*(control)=0.5772, \**P*(DISC1)=0.0336. Object B: n(control/DISC1)=140/170 trials, main effect of Interaction *F*_1,308_=10.78, \*\**P*=0.0011, *P*(control)=0.9183, \*\*\*\**P*(DISC1)<0.0001). Test (Old object: n(control/DISC1)=81/81 trials, main effect of Interaction *F*_1,160_=2.756, *P*=0.0988, P(control)=0.9833, \**P*(DISC1)=0.0377. Novel object: main effect of Interaction *F*_1,234_=0.1381, *P*=0.7105, n(control/DISC1)=119/117 trials, *P*(control)=0.3006, *P*(DISC1)=0.3896). All trials were from 7-8 mice per group. Dashed lines indicate onset of the events. \**P*<0.05, \*\**P*< 0.01, \*\*\**P*<0.001, \*\*\*\**P*<0.0001. Values represent mean ± SEM.

However, during the NPR test, control mice exhibited a significant elevation of Ca^2+^ activity in CA3 prior to exploration of objects/locations during familiarization. In contrast, DISC1 mice failed to show such Ca^2+^ responses in CA3 prior to exploration (Fig. 3J). Supporting these findings, control mice showed robust reliability of peri-event associated Ca^2+^ dynamics as compared to DISC1 mice (below threshold) in CA3 during familiarization (Fig. 3F). Furthermore, DISC1 mice exhibited a significant Ca^2+^ elevation in CA1 prior to exploration of the objects/locations, as compared to control mice that did not show significant Ca^2+^ elevation during exploration (Fig. 3L). Curiously, DISC1 mice showed abnormally high reliability of perievent associated Ca^2+^ dynamics comparing control mice in CA1 during familiarization (Fig. 3H).

In contrast, during the test phase, no significant alterations of Ca^2+^ activity in CA3 and CA1 were observed between control and DISC1 mice (Fig. 3K, M). Consistently, no significant changes in the reliability of peri-event associated Ca^2+^ dynamics were observed in CA3 or CA1 of control and DISC1 mice during the test (Fig. 3G, I). These data together suggest that a small number of dysregulated newborn neurons induce aberrant activity in CA3 and CA1 during the familiarization (but not the test) phase of the spatial memory process.

### Dysregulated adult-born immature neurons disturb functional connectivity during resting state and inter-regional synchrony during spatial memory behavior

Our data suggest that dysregulated DG newborn neurons are capable of inducing aberrant activity across multiple discrete brain regions, including CA3, CA1, MDTH, and IC. Aberrant activity in these distinct brain regions may result from or lead to aberrant synchrony among these brain regions. It has been well established that the degree of correlated neuronal activity (synchrony) reflects the functional state of networks and circuits (Jasper, 1936). Moreover, ample evidence has implicated that inter-regional synchrony is critical for maintaining normal cognitive functions (Mathalon and Sohal, 2015). Therefore, we sought to investigate whether dysregulated newborn neurons contribute to aberrant inter-regional synchrony. To address this question, we first performed partial correlation analysis to analyze brain-wide network correlation using the dataset obtained from rsfMRI. We chose the ROIs for partial correlation analysis based on our retrograde tracing data (SFig. 4A). Partial correlation analysis confirmed reduced correlation between DG and IC. Moreover, we found reduced correlation between TH and CA1, and increased correlation between CA3 and TH (Fig. 4A-C, SFig. 4C-D). We then generated partial correlation networks and found that the functional connectivity density of DISC1 mice was significantly decreased (26% less compared to control mice) (Fig. 4D-F). Consistent results were achieved with different thresholding criteria (SFig. 4B). The degree of individual nodes distributed significantly different in control and DISC1 networks, and there were much fewer high-degree nodes in the DISC1 network, indicating that the functional network connectivity was re-organized with altered hub nodes (Fig. 4G-H). Furthermore, we measured the node weight change of each ROI, and found TH, CA3, and CA1 as the most altered brain regions in DISC1 mice (Fig. 4I). These results support the involvement of DG, CA3, CA1, TH, and IC as a neural network disrupted by dysregulated newborn neurons.

**Figure 4.**
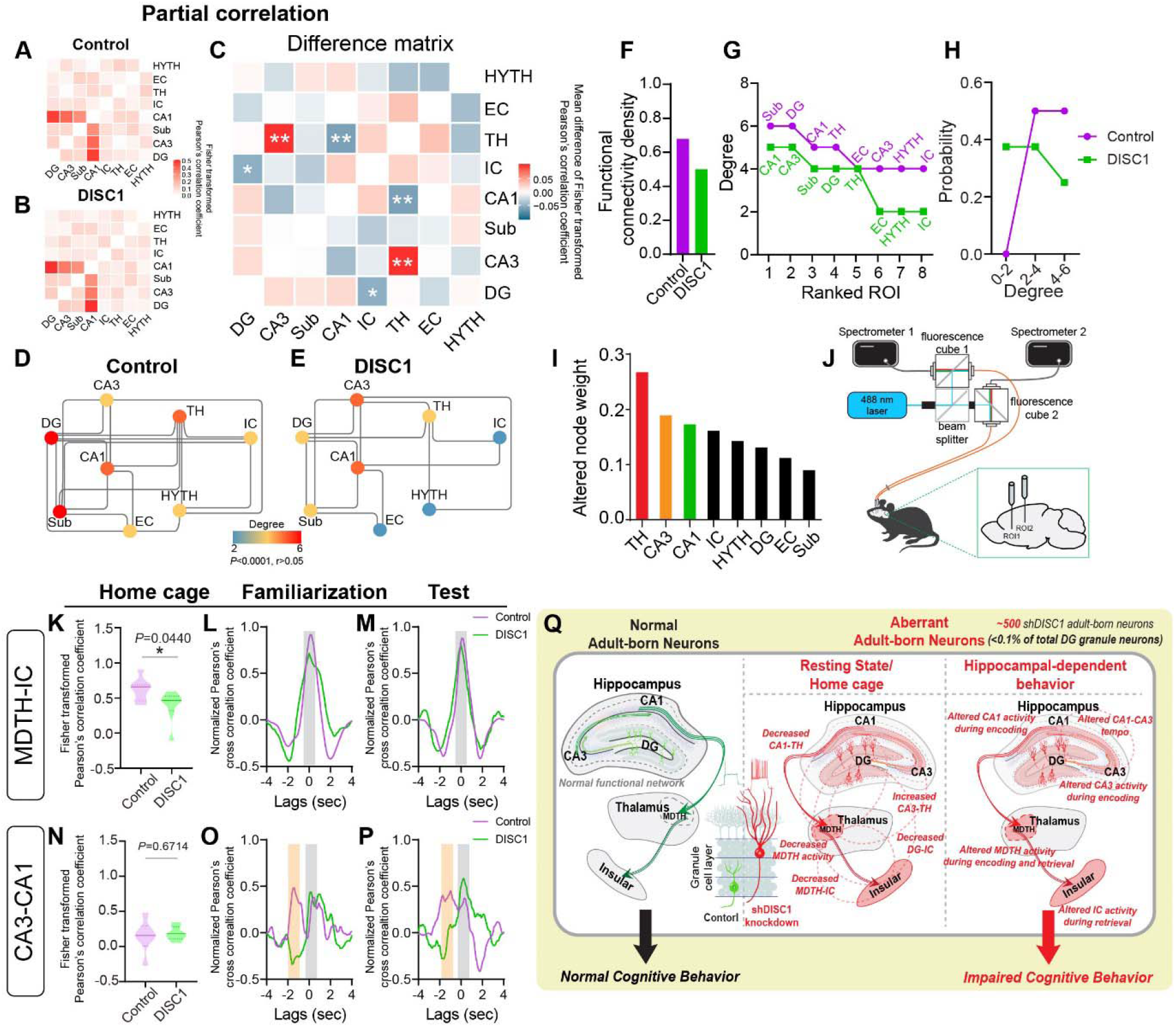
Dysregulated adult-born immature neurons disturb functional connectivity during resting state and inter-regional synchrony during spatial memory behavior. (A-B) Partial correlation matrix. (C) Mean difference matrix of partial correlation (DISC1-control). Two-way ANOVA test showed the main effect of group: \**P*(DG-IC)=0.0413, *FDR*=0.3852; \*\**P*(CA3-TH)=0.0033, *FDR*=0.0918; \*\**P*(CA1-TH)=0.0071, *FDR*=0.0996. n=8 mice per group. *FDR* were corrected by Benjamini & Hochberg adjustment. (D-E) Summarized partial networks for control and DISC1. n=8 mice per group. (F) Comparison of functional connectivity density of the summarized partial networks (D-E). (G) Ranked node degree distribution. (H) Probability of node degree. (I) Ranked index of altered weight for each individual ROI in partial correlation analysis. (J) Illustration of *in vivo* multi-fiber photometry recording system. (K) Correlation measured by raw signal of MDTH-IC at homecage. \**P*=0.0440, n=7/8 mice for control/DISC1. Two-tailed unpaired Student’s *t* test. (L-M) Averaged time lags between IC to MDTH measured by cross correlation. (N) Correlation measured by raw signal of CA3-CA1 at homecage. *P*=0.6714, n=6/7 for control/DISC1. Two-tailed unpaired Student’s *t* test. (O-P) Averaged time lags between CA1 to CA3 measured by cross correlation. Detailed in SFig. 4. Grey areas indicate shared correlation peaks, orange areas indicate correlation peaks occurred in control but not DISC1. (Q) Model of summarized data. \**P*<0.05, \*\**P*< 0.01.

To further investigate the regional correlation during the spatial memory process, we employed a customized multi-fiber photometry system to simultaneously record Ca^2+^ activity across multiple brain regions in freely moving animals (Fig. 4J). Due to the technical limitation for implanting the optical fibers in CA3, CA1, TH, and IC all at the same time, we managed to implant two optic fibers at a time to simultaneously record MDTH and IC, as well as CA3 and CA1. Interestingly, we found that DISC1 mice exhibit a significant decrease in the correlation of Ca^2+^ signals between MDTH and IC, but not between CA3 and CA1, in the home cage (Fig. 4K, N). Furthermore, during the NPR test, no significant lags were found from IC to MDTH by cross correlation analysis, and the peak of the correlation was around 0 second (Fig. 4L, M, SFig. 4E-F, I-J). It is worth noting that there is an additional correlation peak at around −2~-1 seconds lag time between CA1 to CA3 in control mice during the NPR test (Fig. 4O, P orange areas, SFig. 4G-H, K-L), suggesting that CA3 acts earlier than CA1 during exploration of objects/locations under the normal condition. In contrast, the lag of CA1 to CA3 in DISC1 mice was absent, thus leading to increased CA3-CA1 correlation at lag 0 second, and negative correlation at lag −2~-1 second in DISC1 mice (Fig. 4O, P). These results indicate that dysregulated newborn neurons induce disrupted tempo of inter-regional correlation between CA3 and CA1 during the spatial memory process, and decreased inter-regional correlation between MDTH and IC at baseline.

## Discussion

In this study, using a combination of fMRI, rabies-based retrograde tracing, *in vivo* multifiber photometry recording, and network analysis, we provided the first network-level evidence that a small number of dysregulated adult-born immature neurons (<0.1% of total DG granule neurons) are sufficient to induce brain-wide maladaptation across multiple anatomically discrete regions during the spatial memory process, ranging from local hippocampal regions to distal thalamic and cortical regions. Specifically, we found that a small number of dysregulated adult-born neurons lead to: 1) aberrant activity in CA3 and CA1 during spatial memory encoding along with aberrant CA3-CA1 synchrony during both spatial encoding and retrieval; 2) aberrant MDTH activity at baseline and during both spatial encoding and retrieval; 3) aberrant IC activity during spatial memory retrieval; 4) decreased synchrony between MDTH and IC at baseline; and 5) decreased functional network connectivity density and shifted network hubs at resting state (see the summary model in Fig. 4Q). Therefore, the combination of aberrant activity and inter-regional synchrony within this network induced by dysregulated new neurons collectively contribute to the spatial memory deficits.

A question remaining is how this small population of dysregulated neurons can impact the global network and contribute to the cognitive deficits observed in DISC1 mice. We speculate that the hyperexcitability associated with 18 dpi DISC1-deficient newborn neurons may contribute to aberrant activity and functional connectivity across multiple brain regions involved in spatial memory. Supporting this view, our preliminary data showed that some network abnormalities associated with hyperexcitability of dysregulated newborn neurons, such as reduced MDTH activity and MDTH-IC correlation, were abolished when these DISC1-deficient neurons reached a later developmental stage and became hypoexcitable (SFig. 5). These findings support a potential stage-dependent contribution of dysregulated newborn neurons with unique physiological properties in regulating the output connectivity of brain-wide functional network for spatial memory. Recent computational modeling has shown that adding a small number of active newborn neurons (~0.2% of dentate neuron population) to the DG network can have a profound effect on memory function (Tran et al., 2019), further supporting that small changes in hippocampal neurogenesis are sufficient to profoundly impact cognitive functions.

Our studies suggest that adult-born neurons with unique properties can serve as an active modulator of local and distal circuitry to shape mature neuron firing, synchronization, and network connectivity. The involvement of distal brain regions (MDTH and IC) in this spatial memory network mediated by dysregulated adult-born immature neurons is unexpected, because these two brain regions have not previously been implicated in hippocampal dependent spatial memory, though both of them have been implicated in cognitive functions (Gogolla, 2017; Parnaudeau et al., 2013). In addition, we found that MDTH and IC are not only highly anatomically connected, but also highly correlated in their Ca^2+^ signals at home cage and during spatial memory. Synchronization of two discrete brain regions has been thought to be critical for behavior-dependent functional coupling of neural circuits through spike-time coordination (Ito et al., 2018). Interestingly, DISC1 mice exhibited decreased correlation between MDTH and IC at home cage, but did not alter their correlation during spatial memory task. Given that both MDTH and IC activities in DISC1 mice are altered during spatial memory, we speculate that hyperexcitable DISC1-deficient immature neurons may promote maladaptive states of brain-wide neural network and impair behavior by synchronizing discrete brain regions with aberrant activity. It remains to be determined whether MDTH and IC represent the common downstream targets of adult-born immature neurons with enhanced excitability, or they are unique targets of hyperexcitable DISC1-deficient adult-born immature neurons.

Recent conflicting reports about whether adult hippocampal neurogenesis occurs in humans raise questions about its significance for human health and the relevance of animal models. Drawing upon published data, the field now generally agreed that low-level hippocampal neurogenesis persists in adult humans throughout aging (Boldrini et al., 2018; Snyder, 2019; Spalding et al., 2013). Given the low-level human hippocampal neurogenesis in adulthood, our studies showing 500 deficient neurons can impact brain-wide network have broad implications for the functional role of human hippocampal neurogenesis.

## Acknowledgements

We thank the members of Song and Shih labs for their helpful inputs on this project. We thank Bentley R. Midkiff in the UNC Translational Pathology Laboratory (TPL) for expert technical assistance. The UNC Translational Pathology Laboratory is supported in part, by grants from the NCI (5P30CA016086-42), NIH (U54-CA156733), NIEHS (5 P30 ES010126-17), UCRF, and NCBT (2015-IDG-1007). Z.H. is supported by NIH (U01 EB023686 and U41 HG0077346), and Tata Consultancy Services. We thank Joseph R Merrill from BRIC (UNC Biomedical Research Imaging Center) for fMRI scanning. This work was supported by grants awarded to J.S. from NIH (R01MH111773-01, R01MH122692-01, R21AG058160, R21NS104530).

## Author contributions

H.B. and J.S. designed the experiments, interpreted the results and wrote the paper. H.B. performed all the experiments and analysis. Z.H., S.L., W.B. and Y.I.S. contributed to fMRI experiments. R.K. contributed to behavior experiments. T.C. and Y.I.S. contributed to fiber photometry experiments. Y.L. contributed to electrophysiology. H.A.S and I.W. provided rabies tracing materials. Y.Y. assisted fiber photometry and network data analysis. Z.H. and S.E.B. assisted with statistics (SEB has not reviewed the final manuscript due to his medical leave). S. G-A., Z.R.L, and J.H. provided retrovirus.

## STAR Methods

### KEY RESOURCES TABLE

| REAGENT or RESOURCE | SOURCE | IDENTIFIER |
| --- | --- | --- |
| Antibodies |  |  |
| Goat anti GFP | Rockland | Cat# 600-101-215;<br>RRID: AB_218182 |
| Rat anti mCherry | Life Tech | Cat# M11217;<br>RRID:<br>AB_2536611 |
| Bacterial and Virus Strains |  |  |
| AAV8/DJ-CaMKII-GCaMP6f | University of Washington | Dr. Garret Stuber |
| AAV5-CaMKII-GCaMP6f-WPRE-SV40 | Addgene | #100834-AAV5 |
| AAV5-CaMKII-GFP | UNC Vector Core | N/A |
| AAV5-CaMKII-Cre | UNC Vector Core | N/A |
| AAV5-CAG-tdTomato | UNC Vector Core | N/A |
| Retrovirus-Ubi-GFP-WPRE | University of Texas at San Antonio | Dr. Jenny Hsieh |
| Retrovirus-Ubi-shDISC1-GFP-WPRE | University of Texas at San Antonio | Dr. Jenny Hsieh |
| AAV-Syn-FLEX-splitTVA-eGFP-tTA | Massachusetts Institute of Technology | Dr. Ian R. Wickersham |
| AAV-TREtight-BFP2-B19G | Massachusetts<br>Institute of<br>Technology | Dr. Ian R.<br>Wickersham |
| RVΔG-4mCherry (EnvA) | Massachusetts<br>Institute of<br>Technology | Dr. Ian R.<br>Wickersham |
| Biological Samples |  |  |
| Chemicals, Peptides, and Recombinant Proteins |  |  |
| Critical Commercial Assays |  |  |
| Deposited Data |  |  |
| Experimental Models: Cell Lines |  |  |
| Experimental Models: Organisms/Strains |  |  |
| Mouse: C57BL/6J | Jackson Laboratory | 000664 |
| Oligonucleotides |  |  |
| Recombinant DNA |  |  |
| Software and Algorithms |  |  |
| ImageJ(Fiji) | https://imagej.net/ | RRID:SCR_003070 |
| FV10-ASW | Olympus | RRID:SCR_014215 |
| Aperio ScanScope FL | Leica Biosystems | N/A |
| Versa Leica Biosystems | Leica Biosystems | N/A |
| Paravision | Bruker | RRID:SCR_001964 |
| ITK-SNAP | http://www.itksnap.org/ | RRID:SCR_002010 |
| pClamp 10 | Axon | RRID:SCR_011323 |
| Clampfit | Axon | N/A |
| OceanView | Ocean Optics | N/A |
| MATLAB | MathWorks | RRID:SCR_001622 |
| R | https://www.r-project.org/ | RRID:SCR_001905 |
| R Studio | https://rstudio.com/ | N/A |
| Python | https://www.python.org/ | RRID:SCR_008394 |
| Cytoscape | https://cytoscape.org/ | RRID:SCR_003032 |
| GraphPad Prism | Graphpad | RRID:SCR_002798 |
| Other |  |  |

### RESOURCE AVAILABILITY

#### Lead Contact

Further information and requests for resources and reagents should be directed to and will be fulfilled by the Lead Contact, Juan Song.

#### Materials Availability

This study did not generate new unique reagents.

#### Data and Code Availability

The data that support the findings of this study are available from the corresponding authors J.S. upon reasonable request. Computer codes used to generate results reported in the manuscript are available to editors and referees upon request. Codes will be shared after publication.

### EXPERIMENTAL MODEL AND SUBJECT DETAILS

#### Animals

6-8 week male C57BL/6J (Jackson Laboratory NO: 000664) mice were used for all experiments. All animals were experimentally and drug naive before use. Animals were group housed and bred in a dedicated husbandry facility with 12/12 light-dark cycles with food and water ad libitum and under veterinary supervision. Animals subjected to surgical procedures were moved to a satellite housing facility for recovery with the same light-dark cycle. Surgical strategies were carefully designed to combine both shControl and shDISC1 injected mice in the same cage. All procedures were conducted in accordance with the NIH Guide for the Care and Use of Laboratory Animals, and with approval of the Institutional Animal Care and Use Committee at the University of North Carolina at Chapel Hill (UNC).

### METHOD DETAILS

#### Stereotaxic surgery

Young adult mice (8-10 weeks) were anesthetized under 1%-2% isoflurane in oxygen at 0.6-0.8 LPM flow rate. Injection coordinates were based on “The Mouse Brain in Stereotaxic Coordinates” (Second Edition, Paxinos and Franklin, 2001) and pilot experiments. Virus was injected by microsyringe (Hamilton, 33GA) and microinjection pump (Harvard Apparatus), at the rate of 100 nl/min with the following coordinates. DG (dorsal AP: −2.0 mm, ML: ±1.5 mm, DV: − 2.3 mm; ventral AP: −3.0 mm, ML: ± 2.5 mm, DV: −3.0 mm); CA1 (AP: −2.0 mm, ML: +1.5 mm, DV: −1.45 mm); CA3 (AP: −2.5 mm, ML: +2.8 mm, DV: −3.0); IC (AP: +1.6 mm, ML: +2.9 mm, DV: −3.5 mm); MDTH (AP: −1.65 mm, ML: +0.3 mm, DV: −3.8 mm). For retroviral birth dating, mice were injected with Retrovirus-Ubi-GFP-WPRE or Retrovirus-Ubi-shDISC1-GFP-WPRE to bilateral dorsal and ventral DG, 0.3 μL/site, 4 sites in total. For fiber photometry, mice were first injected with retrovirus to DG as described. Additional 0.3 μL pre-mixed AAV-CamKII-GCaMP6f-WPRE-SV40 / AAV5-CAG-tdTomato (mixed ratio 3:1) were injected to recording site. Optical fibers (Thorlabs, Ø1.25 mm Multimode LC/PC Ceramic Ferrule, Ø230 mm Hole Size) were then implanted 0.2 mm above GCaMP injection site. For monosynaptic rabies tracing from IC or MDTH, animals were first injected with 0.3 μL pre-mixed helper virus AAV-CaMKII-Cre / AAV-Syn-FLEX-splitTVAeGFP-Tta / AAV-TREtight-BFP2-B19G (mixed ratio 1:2.5:2.5) to the right IC. After 7 day, animals had the second injection of 0.3 μL RVΔG-4mCherry (EnvA) to the same injection site. Then animals were transferred to a quarantined cubicle for special housing and monitoring. 7 days post rabies injection, animals were perfused and brain tissues were collected. For monosynaptic rabies tracing from CA1 or CA3, animals were first injected 0.3 μL pre-mixed helper virus AAV-CaMKII-Cre / AAV-Syn-FLEX-splitTVAeGFP-tTA (mixed ratio 1:5) to CA1 or CA3. After 7 day, animals received the second injection of 0.3 μL RVΔG-4mCherry (EnvA) to their right MDTH. Then animals were transferred to a quarantined cubicle for special housing and monitoring. 7 days post rabies injection, animals were perfused and brain tissues were collected. AAV8/DJ-CaMKII-GCaMP6f virus was obtained from Dr. Garret Stuber Lab (University of Washington). AAV5-CamKII-GCaMP6f-WPRE-SV40 was purchased from Addgene (#100834-AAV5). AAV5-CaMKII-GFP, AAV5-CaMKII-Cre, AAV5-CAG-tdTomato were obtained from UNC Vector Core. Virus titer is 2-4×10^12^ vg/mL. Retroviral vectors Retrovirus-Ubi-GFP-WPRE and Retrovirus-Ubi-shDISC1-GFP-WPRE were obtained from Dr. Jenny Hsieh (University of Texas at San Antonio). Monosynaptic tracing related helper virus AAV-Syn-FLEX-splitTVAeGFP-tTA, AAV-TREtight-BFP2-B19G and rabies RVΔG-4mCherry (EnvA) were obtained from Dr. Ian R. Wickersham (Massachusetts Institute of Technology).

#### BOLD-fMRI

##### Imaging

The separate groups of mice were used for resting state fMRI approaches. 18 days post retrovirus injection, the mice were anesthetized and placed into the Bruker 9.4T preclinical MRI scanner. Total 500 volumes (25 min) of the isotropic BOLD signal were acquired for the whole brain using single shot gradient echo EPI sequence (TR: 3000 ms, TE: 10 ms, flip angle: 80 deg, resolution: 0.3 x 0.3 x 0.3 mm^3^, FOV: 19.2mm x 19.2mm x 7.8mm). During the scan, the mice were anesthetized with 0.8 ~ 1 % Isoflurane. The body temperature and the breathing rate were monitored real-time while maintained within a range of 37.0 ± 0.5 and ~120 bpm respectively. The anatomical brain image of each mice was also acquired using turbo RARE sequence at T2 (TR: 1000ms, TE: 11ms, Effective TE: 33, RARE factor 8, resolution: 0.15 x 0.15 x 0.15 mm^3^, FOV: 22.8 x 14.4 x 7.8) in order to determine subject-wise anatomical validation of fMRI results.

##### Preprocessing

The data were manually skull stripped in order to improve the accuracy of brain registration process. Core preprocessing procedures including co-registration, slice timing correction and motion correction were applied for each animal using AFNI toolbox. Subsequently, brain image of each subject was realigned into the mouse brain MRI template using ANTs. Then, six rigid head motion parameters and their derivatives were used as a regressor to filter off the nuisance signal, and a bandpass filter in the 0.01 to 0.1 Hz range was applied to detect low-frequency BOLD oscillation in resting state. Finally, the Gaussian smoothing kernel with full-width half-maximum at 0.4 mm was applied to reduce random noise at the spatial domain.

#### Animal behaviors

Animals with virus injection and optical fiber implantation were allowed to recover for 2 weeks. Three days before the behavior test, animals were transferred to the behavior room daily and handled carefully by the experimenter to adapt to the environmental stress. The behavior chamber was exposure to the animals for habituation. On recording day, animals were connected to the fiber patch chord, and recorded baseline activity for 20 mins. In novel place test, test animals were first placed into the chamber with two identical objects and familiarized with the environment for 10 mins. Then animals were moved to their home cage. One object (Old location) stayed in the same position, while the other object (Novel location) was moved to a new position. 5 mins later, same animals were then reintroduced to the chamber for 10 mins test. The whole process was recorded by the video camera on top of the chamber.

#### Fiber photometry

For simultaneous dual-channel fiber-photometry, a single 488 nm laser beam (OBIS 488LS-60, Coherent, Inc.) was equally split into two laser beams using a beam splitter (CCM1-BS013, Thorlabs, Newton, NJ), and then the two laser beams were respectively launched into two fluorescence cubes (DFM1, Thorlabs, Newton, NJ). An extra neutral density filter (NEK01, Thorlabs, Newton, NJ) was placed in front of the input of the beam splitter for additional adjustment of the laser power. For each channel, a dichroic mirror (ZT488/561rpc, Chroma Technology Corp) inside the fluorescence cube reflected and launched the laser beam through an achromatic fiber port (PAFA-X-4-A, Thorlabs, Newton, NJ) into a multi-mode optical fiber patch cable. The distal end of the patch cable was connected to the animal through an implantable optical fiber probe, serving for both laser delivery and GCaMP6f fluorescence collection. The emission fluorescence traveled back through the patch cable to the fluorescence cube, and passed through the dichroic mirror and an emission filter (ZET488/561 m, Chroma Technology Corp, Bellows Falls, VT), then was launched through an aspheric fiber port (PAF-SMA-11-A, Thorlabs, Newton, NJ) into the core of an AR-coated 200/230 mm core/cladding multi-mode patch cable (M200L02S-A, Thorlabs, Newton, NJ). The AR-coated multi-mode patch cable was connected to a spectrometer (Ocean Optics, Largo, FL) for spectral data acquisition, which was operated by OceanView (Ocean Optics, Largo, FL), an PC software under Windows environment. IC and MDTH recordings were done within the same animals simultaneously. CA3 and CA1 recordings were done within the same animals simultaneously. The *in vivo* correlation data were analyzed by using the same recording dataset respectively.

#### Immunohistochemistry

Mice were anesthetized with ketamine, and perfused with ice-cold 4% paraformaldehyde (PFA) in PBS. Brains were collected and placed in 4% PFA overnight, and switched to 30% sucrose for 2-3 days until they were fully submerged. Brains were sectioned on a microtome at the thickness of 40 μm, and stored in the anti-freeze solution at 20°C for further usage. For staining, brain sections were first mounted on charged slides and washed twice with TBS, followed by permeabilization with 0.5% Triton-100 TBS for 20 minutes. After washed with TBS+ (TBS with 0.05% Triton-100) for 5 minutes and blocked with 3.5% donkey serum in TBS+ for 30 minutes, sections were incubated in the primary antibody overnight at 4°C on the shaker. On the second day, brain sections were washed with TBS+, and then incubated with secondary antibody at room temperature for 2 hours. Sections were counterstained with DAPI for 10 minutes, and covered by mounting media. Primary antibodies: Anti-GFP (goat, 1:500, Rockland), Anti-mCherry (rat, 1:500, Life Tech).

#### Imaging

Confocal images were acquired by Olympus FLUOVIEW1000 confocal microscopy, under 40x Oil (NA1.30), XY-resolution 0.4975 mm per pixel, Z-resolution 1.0 or 1.5 mm per slice. Tiled images were acquired based on locations of the fluorescent signals, and images were stitched after acquisition using the Olympus FLUOVIEW imaging software. Brightness and contrast were adjusted in ImageJ.

Monosynaptic rabies input screening images were acquired and stitched by UNC Translational Pathology Laboratory. The process of quantitative image analysis begins with the acquisition of high-resolution digital slides. Frozen sections of mouse brain tissue were scanned on either an Aperio ScanScope FL or the Versa (Leica Biosystems) using a 20x objective. Images were then uploaded to eSlide Manager and visualized with ImageScope 12.3.3 (Leica Biosystems). Files from the ScanScope FL are in Aperio Fused Image (.afi) format, and files from the Versa are in Leica SCN format. The image resolution is 0.46290 microns per pixel, and the image depth is 16 bits per channel for the ScanScope FL scans. The image resolution is 0.32182 microns per pixel, and the image depth is 8 bits per channel for the Versa scans.

#### Electrophysiology

Animals were anesthetized with 2% isoflurane and then decapitated. The brain was rapidly removed and immersed in an ice-cold N-Methyl-D-glucamine (NMDG)-containing cutting solution (in mM): 92 NMDG, 30 NaHCO_3_, 25 glucose, 20 HEPES, 10 MgSO_4_, 5 sodium ascorbate, 3 sodium pyruvate, 2.5 KCl, 2 thiourea, 1.25 NaH_2_PO_4_, 0.5 CaCl_2_ (pH 7.3, 310 mOsm). Transverse brain slices (280 μm thickness) containing the hippocampus were cut by using a vibratome (VT1200 S, Leica, USA). After cutting, slices were allowed to recover in the cutting solution saturated with 95% O_2_ and 5% CO_2_ for 8 min at 34°C. The slices were then transferred and incubated in HEPEs holding solution containing (in mM): 92 NaCl, 30 NaHCO_3_, 25 glucose, 20 HEPES, 5 sodium ascorbate, 3 sodium pyruvate, 2.5 KCl, 2 thiourea, 2 MgSO_4_, 2 CaCl_2_, 1.25 NaH_2_PO_4_ (pH 7.3, 310 mOsm) at room temperature for at least 1 hour before recording.

Whole-cell patch-clamp recordings were made from GFP positive neurons visualized under fluorescence and infrared differential interference contrast microscopy on an upright BX51WI microscope (Olympus, Japan). Slices were placed in a perfusion chamber and submerged in continuously flowing oxygenated recording artificial CSF containing the following (in mM): 125 NaCl, 25 NaHCO_3_, 25 glucose, 2.5 KCl, 1.25 NaH_2_PO_4_, 2 CaCl_2_, and 1 MgCl_2_ (pH 7.3, 310 mOsm). Electrophysiological recordings were made with a Multiclamp 700B amplifier and digitized with a Digidata 1440A (Axon Instruments, USA). Patch electrodes were made by pulling glass capillaries (World Precision Instruments, USA) on a Puller (PC-10, Narishige, Japan). The pipette resistance was typically 4-6 MQ.

### QUANTIFICATION AND STATISTICAL ANALYSIS

#### BOLD-fMRI Analysis

##### Correlation analysis

In order to detect brain regions affected by DG, we first calculated the pairwise correlation between bilateral DG (biDG) and IC, and then compared pairwise correlation between and the control and DISC1 mice. Specifically, correlations between biDG and IC (e.g. left IC) was calculated with Pearson correlation. The difference of each pairwise correlation between control and DISC1 mice was tested as follows: (1) all correlation coefficients were first normalized using Fisher z-transformation method; (2) for correlations between DG and regions with left and right sub regions, a p-value was calculated with a linear model with the positioning information (“left” or “right) as a covariate.

##### Partial correlation analysis

ROIs were manually defined in the mouse brain atlas file, including Insular cortex (IC), dentate gyrus (DG), subiculum (Sub), medial dorsal thalamus (MDTH), paracentral thalamic nucleus (PCN), hypothalamus (HYTH), entorhinal cortex (EC). Thalamus (TH) shown in Fig. 4 is merged by MDTH and PCN. The ROIs were further separated into left and right sub regions. The time-series signals of each sub brain region were calculated as the average of all within-region voxels. For each of the sixteen mice (8 control and 8 DISC1), a connectivity network of the predefined brain regions was constructed. Specifically, the connectivity was inferred by partial correlations controlling for all the other brain regions as well as their positioning information (*P*<0.05 and partial correlation coefficient >0.05). For connectivity between each pair of brain regions in at least one representative network, we test their difference between DISC1 mice and the control mice. Specifically, the partial correlation coefficients were normalized using Fisher z-transformation method and the significance was detected using unpaired *t*-tests followed by Benjamini & Hochberg adjustment. Next, two representative networks (DISC1 and control networks) were generated by summarizing the individual networks: two brain regions were considered as connected if (1) the summary p-value calculated by Fisher’s combined probability test was less than 0.0001; and (2) the average partial correlation coefficient was higher than 0.05. The networks were visualized by Cytoscape. The altered node weight was measured as the sum of absolute changes of partial correlation coefficients of each individual brain regions. The functional connectivity density and node degree were measured with build-in tool in Cytoscape. Both correlation and partial correlation analysis were done with the same dataset in R.

#### Behavior Analysis

The animal behavior analysis was scored by experienced analyst unbiasedly. Time of exploring and frequency was measured manually based on video. Locomotion of NPR test were measured by ImageJ. Peri-event analysis was based on sniffing event for either object/location. Time courses of behavioral video and fiber photometry recording data were synchronized to the start time point.

#### Fiber Photometry

##### Unmixing and Ca^2+^ activity analysis

Fiber photometry data were acquired in a spectrum format (348-1136 nm) at 10 Hz sampling rate (Fig. 1J). Customized scripts were applied to process and analyze the data matrix in MATLAB R2017a. Mixed spectrum acquired by fiberphotometry was analyzed using a spectral linear unmixing algorithm. At time point t, the mixed spectrum Y(t) is model as:

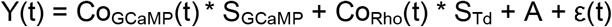

Where S_GCaMP_ and S_Td_ are the normalized reference emission spectra of GCaMP and tdTomato respectively. Co_GCaMP_(t) and Co_Td_(t) are the unknown regression coefficients corresponding to the GCaMP and tdTomato signals respectively. A is the unknown constant, and ε(t) is random error. Next, the Co_GCaMP_(t) was detrended via high-passed filtering with cut-off frequency at 0.01 Hz, and then was further motion-corrected and normalized to fractional fluorescence changes (ΔF/F) using the following equation:

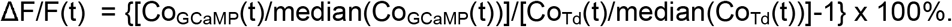

For the following quantifications, high Ca^2+^ activity (Ca^2+^_high_) was determined by thresholding the ΔF/F(t) at 3 x standard deviation (SD) of ΔF/F(t):

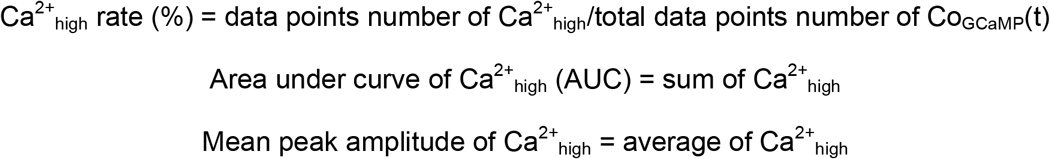

Behavior-related neuronal activities were extract by perievent analysis, and the results are presented in z-score. Each animal was validated by their signal quality. Misplacements and poor signals were excluded from analysis.

##### Power frequency analysis

The time-frequency energy distributions of the GCaMP responses were analyzed by continuous Morlet wavelet transform using homemade MATLAB scripts. The mean power spectrums were the average of energy distributions across the period during home cage recording, or during NPR events.

##### Peri-event analysis

Peri-event analysis of NPR test were based on sniff timing of the object. Events were then extracted (1 sec before and 0.5 sec after) from the preprocessed data and normalized to z-score.

##### Calcium response reliability

The cross covariance was calculated between all pair-wise combinations of calcium response trials (within stimulus size) to every coding event with time bin of 0.1 sec. The covariance function estimates the mean-removed cross-correlation between the two sequences of random processes, thus avoiding the contribution of the mean response intensity to the final calculated reliability. The resulting cross-covariance values at zero-lag (normalized by the average autocovariance function) for each trial were used to quantify the Ca^2+^ timing reliability for each event statistically. Randomly selected non-event Ca^2+^ signals from the same recordings were used as negative control respectively. Robust reliability was thresholding at 0.01.

##### Correlation analysis for multi-fiber photometry

Pearson’s correlation analysis was applied for multi-fiber photometry recording data with “corr” function with homemade MATLAB scripts. Time lags of NPR events were measured by “xcorr” function with homemade MATLAB scripts. Peri-event trials were shuffled for random combinations. Each time, ¾ of the trials were selected and averaged for Pearson’s cross correlation measure. Time lags from 50 different shuffled combinations were then averaged for data presentation.

#### Monosynaptic Rabies Tracing

Annotated regions were drawn around appropriate anatomical features on each image. The annotated FL scans were then analyzed with the Aperio Color Deconvolution v9 algorithm configured as a custom macro for this assay. Annotated Versa scans were analyzed with Definiens Architect XD 2.7, Build 60765 x64 with Tissue Studio 4.4.2 (IF Portal). To create a customized analysis macro with the Color Deconvolution v9 algorithm, the mCherry intensity threshold was set to distinguish negative and positive pixels. Data output included the Area of Positive Pixels as well as Percent Positive. For Definiens Tissue Studio, the annotated regions were imported into the analysis program to calculate marker area. Then, the intensity threshold was set for the mCherry detection. The analysis output included all quantitative results as well as color-coded overlays that represented marker intensity.

#### Electrophysiology

For Intrinsic membrane property recording, GFP+ cells in the DG were recorded in current-clamp mode using a K-gluconate-based intracellular solution (in mM): 130 K-gluconate, 20 HEPES, 4 MgCl_2_, 4 Na-ATP, 2 NaCl, 0.5 EGTA, 0.4 Na-GTP (pH 7.25, 290 mOsm). Relations between firing frequency and injected current were examined by measuring action potentials elicited by somatic current injections (500-ms duration) from 0 to 240□pA with 20 pA step-wise increments. All data were acquired at a sampling frequency of 10 kHz, filtered at 2 kHz and analyzed using pClamp10 software (Molecular Devices). Access resistance of recordings was <30 MQ and monitored throughout the experiment. Data were discarded if the access resistance changed by >20%.

#### Statistical Analysis

Mouse identity and experimental manipulation was coded to allow blinding for all experiments. Sample sizes were based on previous publications with extra samples to account for potential excluded data. Mice with off-target viral injections or expression outside the area of interest were excluded from analysis after post hoc examination of fluorescence expression. Sample sizes was represented by ‘n’ for animal number or cell number and reported in figure legends. Values represent mean ± SEM in bar or superimposed symbols/line plots. Solid and dashed lines represent median and quartiles in violin plots. All statistics data were analyzed in GraphPad Prism8, R or Python. Shapiro-Wilk test was performed for normality. Two-tailed Wilcoxon signed rank test, two-tailed paired/unpaired Student’s *t* test, and two-way repeated measured ANOVA followed by Sidak’s post hoc test, were performed for different dataset. *P* values from multiple comparisons were corrected with Benjamini & Hochberg adjustment and reported as *FDR*. Fisher’s combined probability test were performed for the representative partial networks. Detailed statistics methods are described in the figure legends.

---

**Supplementary Figure 1 (related to Figure 1).**
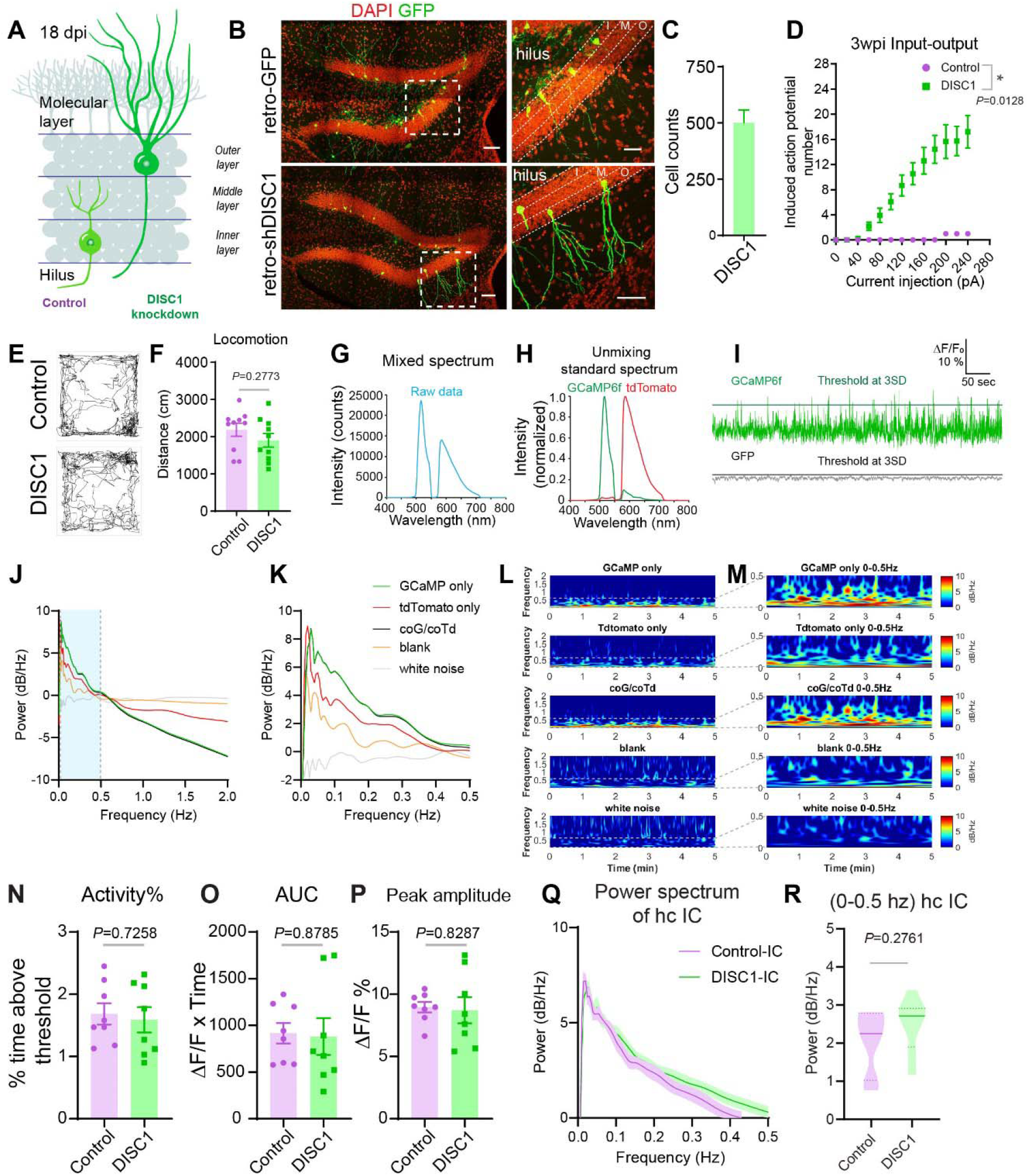
Characterization of DISC1 knockdown phenotypes and *in vivo* Ca^2+^ fiber photometry recording in IC. (A) Summary illustration of developmental deficits in *DISC1*-knockdown newborn neurons. (B) Composite confocal images showing morphological deficits of *DISC1*-knockdown newborn neurons. I, M, O indicate inner, middle, and outer granule cell layers, respectively. Scale bar, 100 μm (left), right 50 μm (right zoomed in). (C) Quantification of retro-shDISC1 labeled cells per DG (n=2). (D) Input-output relationship between injected current and firing frequency of control and shDISC1 retroviral labeled neurons. (n=2/13 cells for control/DISC1. Mixed effects analysis: main effect of Interaction, *F*_12,144_=5.806, \*\*\*\**P*<0.0001. Main effect of Group, *F*_1.311,15.72_=6.975, \**P*=0.0128). (E) Sample locomotion trace of control and DISC1 mice during novel place test. (F) Distance traveled during novel place test. (n= 10/10 for each group, two-tailed unpaired Student’s *t* test, *P*=0.2773). (G) Sample data showing raw mixed spectrum collected from a spectrometer. (H) Standard spectrums of GCaMP6f and tdTomato used for unmixing raw data. (I) Upper: Representative fiber photometry trace of GCaMP6f signal during home cage recording. Lower: Representative GFP signal as negative control with non-neuronal activity. Solid line is the threshold at 3x standard deviation of individual recording. (J) Sample traces of power spectral density of IC during homecage recording with various control conditions. (K) Power spectral density of the dominant frequency component (0-0.5Hz). (L) Morlet wavelet transformed power of IC homecage Ca^2+^ activity in frequency and time domain. (M) Dominant frequency component of IC homecage Ca^2+^ activity in frequency and time domain. Blank indicates recording in the absence of fluorescence reporter. White noise is randomly generated noise signal. (N-P) IC homecage Ca^2+^ activity. (N) The percentage of high Ca^2+^ activity events among total recording session, *P*=0.7258. (O) Area under the curve of high Ca^2+^ activity events, *P*=0.8785. (P) Average peak amplitude of high Ca^2+^ activity events, *P*=0.8287. Two-tailed unpaired Student’s *t* test. n=8 per group. (Q) Power spectrum of IC Ca^2+^ activity at homecage. (R) Comparison of averaged power density of 0-0.5 Hz in IC at homecage. (n=7/8 mice for control/DISC1, two-tailed unpaired Student’s *t* test, *P*=0.2761). Values represent mean ± SEM.

**Supplementary Figure 2 (related to Figure 2).**
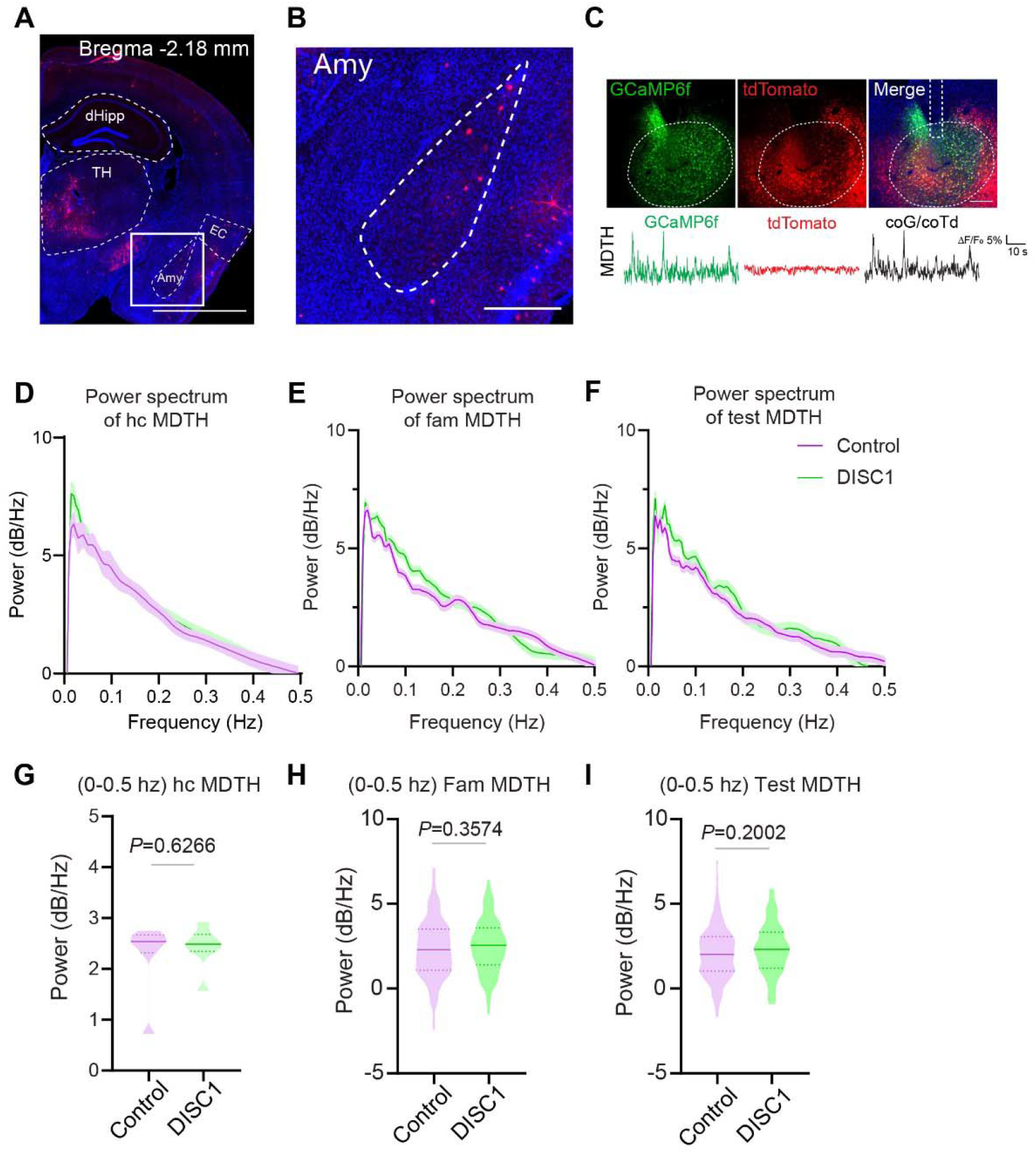
*In vivo* Ca^2+^ fiber photometry recording in MDTH. (A) Images of distal inputs from amygdala (Amy). Scale bar, 2 mm. (B) Zoomed image of Amy inputs. Scale bar, 400 μm. (C) Low-mag sample image showing GCaMP6f and tdTomato viral co-expression and fiber trace in MDTH. Scale bar, 2 mm. (D-F) Power spectral density of MDTH Ca^2+^ activity at homecage, during NPR familiarization and NPR test. (G-I) Comparison of averaged power spectral density of 0-0.5 Hz in MDTH at homecage, during NPR familiarization and NPR test. (n(fam)= 264/214 trials and n(test)= 204/118 trials from 8 mice per group. Two-tailed unpaired Student’s *t* test, *P*(homecage)= 0.6266, *P*(fam)= 0.3574, *P*(test)= 0.2002)

**Supplementary Figure 3 (related to Figure 3).**
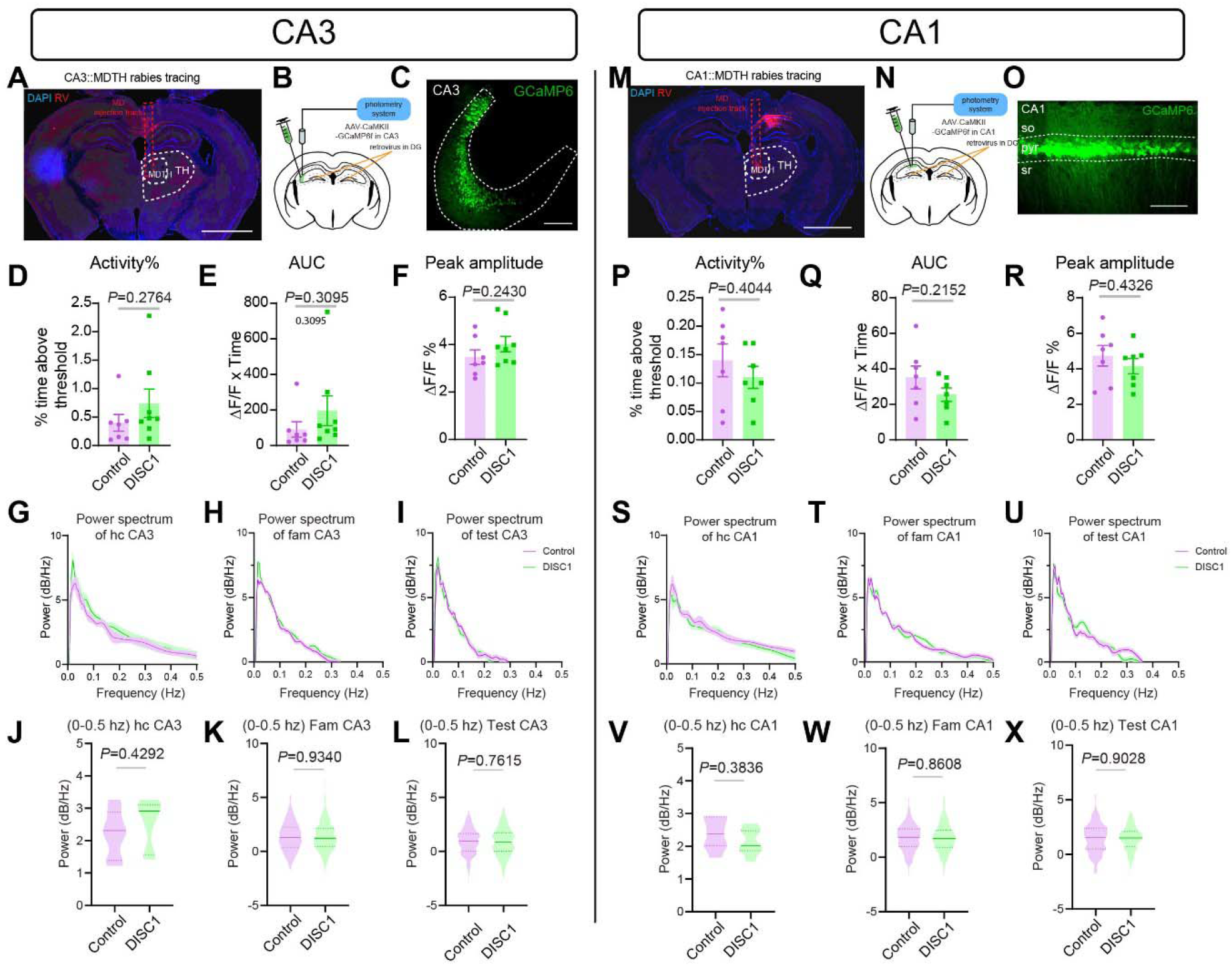
*In vivo* Ca^2+^ fiber photometry recording in CA3 and CA1. (A) Image of rabies injection site to MDTH for CA3::MDTH retrograde tracing. Scale bar, 2 mm. (B) Experimental scheme of fiber photometry recording in CA3. (C) Sample image showing GCaMP6f expression in CA3. Scale bar, 100 μm. (D-F) CA3 homecage Ca^2+^ activity. (D) The percentage of high Ca^2+^ activity events among total recording session, *P*=0.2764 (E) Area under the curve of high Ca^2+^ activity events, *P*=0.3095, (F) Average peak amplitude of high Ca^2+^ activity events, *P*=0.2430. Two-tailed unpaired Student’s *t* test. n=7/8 mice (Control/DISC1). (G-I) Power spectral density of CA3 Ca^2+^ activity at homecage, during NPR familiarization and NPR test. (J-L) Comparison of averaged power spectral density of 0-0.5 Hz in CA3 at homecage, during NPR familiarization and NPR test. (n(homecage)=7/7 mice, n(fam)=303/353 trials and n(test)=200/198 trials from n=7/7 mice for control/DISC1. Two-tailed unpaired Student’s *t* test, *P*(homecage)=0.4292, *P*(fam)=0.9340, *P*(test)=0.7615) (M) Image of rabies injection site to MDTH for CA1::MDTH retrograde tracing. Scale bar, 2 mm. (N) Experimental scheme of fiber photometry recording in CA1. (O) Sample image showing GCaMP6f expression in CA1. Scale bar, 100 μm. (P-R) CA1 homecage Ca^2+^ activity. (P) The percentage of high Ca^2+^ activity events among total recording session, *P*=0.4044 (Q) Area under the curve of high Ca^2+^ activity events, *P*=0.2152 (R) Average peak amplitude of high Ca^2+^ activity events, *P*=0.4326. Two-tailed unpaired Student’s *t* test. n=8 mice per group. (S-U) Power spectral density of CA1 Ca^2+^ activity at homecage, during NPR familiarization and NPR test. (V-X) Comparison of averaged power spectral density of 0-0.5 Hz in CA1 at homecage, during NPR familiarization and NPR test. (n(homecage)=7/7 mice, n(fam)=303/353 trials and n(test)=200/198 trials from n=7/7 mice for control/DISC1. Two-tailed unpaired Student’s *t* test, *P*(homecage)=0.3836, *P*(fam)=0.8608, *P*(test)=0.9028). Values represent mean ± SEM.

**Supplementary Figure 4 (related to Figure 4).**
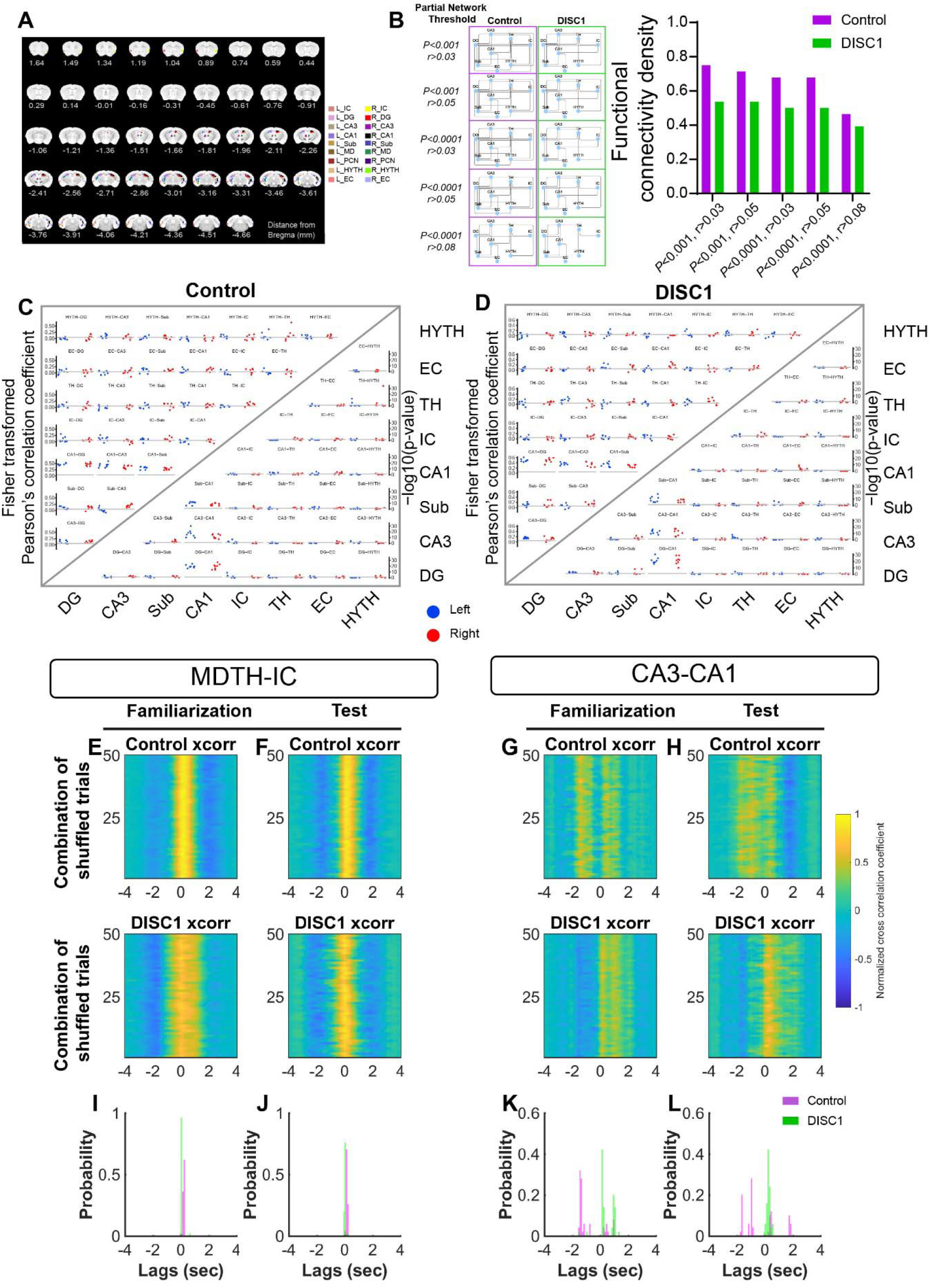
Network analysis based on fMRI and multifiber photometry recording. (A) Defined anatomical ROIs for correlation analysis. Left (L), right (R). Insular cortex (IC), dentate gyrus (DG), subiculum (Sub), medial dorsal thalamus (MDTH), paracentral thalamic nucleus (PCN), hypothalamus (HYTH), entorhinal cortex (EC). Thalamus (TH) shown in figure 4 is merged by MDTH and PCN. (B) Validation of functional network density based on different thresholding. *P* is measured by Fisher’s combined probability test within group, 8 mice per group. (C) Partial correlation coefficient and p-value matrix of control group. (n=8 mice per group). (D) Partial correlation coefficient and p-value matrix of DISC1 group. (n=8 mice per group). Data points represent individual animals. The grey horizontal line is at 0.05 in the correlation triangular matrix, and at 1.30 (=-log10(0.05)) in the p-value triangular matrix. (E-H) Peri-event trials from the same session (eg. MDTH-IC familiarization of control mice) were shuffled for random combinations. Heatmaps showing Pearson’s cross correlation results for 50 random different combinations. (I-L) Corresponding probability histogram of lag time of the “peak of cross correlation”.

**Supplementary Figure 5 (related to discussion).**
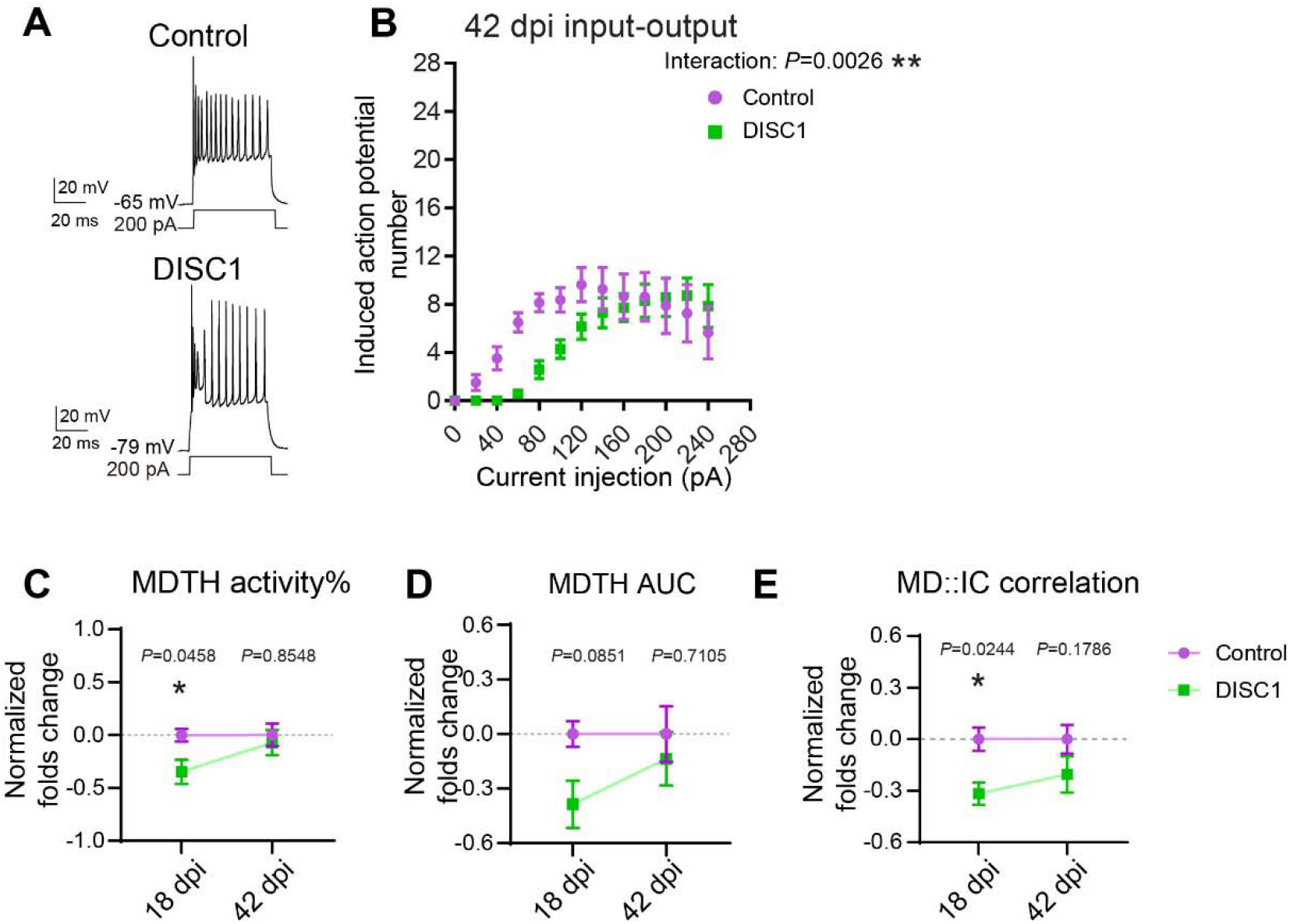
DISC1 phenotypes were abolished at 42 dpi. (A) Sample voltage responses of control (top) and DISC1 (bottom) neurons responding to a 500-ms current injection at 42 dpi. (B) Input-output relationship between injected current and firing frequency of control and shDISC1 retroviral labeled neurons. (n=8/7 cells for control/DISC1. Two-way RM ANOVA: main effect of Interaction, *F*_12,156_=2.686, \*\**P*=0.0026. Main effect of Group, *F*_1,13_=2.027, *P*=0.1781). (C) Normalized comparison of high Ca^2+^ activity events in MDTH at 18 and 42 dpi. (n=8/8 mice for control/DISC1. Two-way ANOVA followed by Sidak’s post hoc: \**P*(18 dpi control vs DISC1)=0.0458*, *P*(42 dpi control vs DISC1)=0.8548). (D) Normalized comparison of area under the curve of high Ca^2+^ activity event in MDTH at 18 and 42 dpi. (n=8/8 mice for control/DISC1. Two-way ANOVA followed by Sidak’s post hoc: *P*(18 dpi control vs DISC1)=0.0851, *P*(42 dpi control vs DISC1)=0.7105) (E) Normalized comparison of MDTH::IC correlation at 18 and 42 dpi. (n=8/8 mice for control/DISC1. Two-way ANOVA followed by Sidak’s *post hoc*: \**P*(18 dpi control vs DISC1)=0.0244*, *P*(42 dpi control vs DISC1)=0.1786).

